# Identification of endothelial-to-mesenchymal transition gene signatures in single-cell transcriptomics of human atherosclerotic tissue

**DOI:** 10.1101/2023.07.18.549599

**Authors:** Lotte Slenders, Marian Wesseling, Arjan Boltjes, Daniek M.C. Kapteijn, Marie A.C. Depuydt, Koen Prange, Noortje A.M. van den Dungen, Ernest Diez Benavente, Dominique P.V. de Kleijn, Gert J. de Borst, Hester M. den Ruijter, Gary K. Owens, Michal Mokry, Gerard Pasterkamp

**Affiliations:** Central Diagnostics Laboratory, Department of Laboratories, Pharmacy and Biomedical Genetics, University Medical Center Utrecht, University Utrecht, Utrecht, The Netherlands; Laboratory of Experimental Cardiology, Department of Cardiology, University Medical Center Utrecht, University Utrecht, Utrecht, The Netherlands; Leiden Academic Centre for Drug Research, Division of BioTherapeutics, Leiden University, Leiden, The Netherlands; Amsterdam University Medical Centers – location AMC, University of Amsterdam, Experimental Vascular Biology, Department of Medical Biochemistry, Amsterdam Cardiovascular Sciences, Amsterdam, The Netherlands; Department of vascular surgery, University Medical Center Utrecht, University Utrecht, Utrecht, The Netherlands; Robert M. Berne Cardiovascular Research Center, University of Virginia, Charlottesville, USA; Department of Molecular Physiology and Biological Physics, University of Virginia, Charlottesville, USA

**Author notes:** Corresponding Author: Prof. dr. Gerard Pasterkamp Central Diagnostics Laboratory, Division Laboratories, Pharmacy and Biomedical genetics University Medical Centre Utrecht, Heidelberglaan 100 3584CX Utrecht The Netherlands. Authors contributed equally to this work (Lotte Slenders, Marian Wesseling).

**Keywords:** EndoMT, Atherosclerosis, scRNA-seq, gene signature

## Abstract

**Rationale:** Endothelial cells can differentiate into mesenchymal-like cells via endothelial to mesenchymal transition (EndoMT). In murine models, cell transitions of EndoMT have been assessed with lineage tracing techniques. Knowledge on molecular mechanisms of EndoMT in human vascular lesions is scarce as studies in human atherosclerosis are limited by observational study designs such as histo-pathological studies.

**Objective:** We aim to identify a human EndoMT gene expression signature by combining experimentally induced *in vitro* EndoMT with lineage-traced pathways from atherosclerotic mice and extrapolate this to human plaque scRNA-seq data.

**Methods and results:** First, we stimulated human coronary artery endothelial cells (HCAEC) with TNFα and TFGβ to trigger EndoMT. We executed transcriptomic analyses and defined multiple temporal patterns of gene expression changes during EndoMT. We used Cdh5-Cre^ERT2^ Rosa-eYFP apoE^-/-^ lineage traced mouse scRNA-seq data to demonstrate that the temporal *in vitro* gene expression changes are reflected in EndoMT trajectories in mice plaque tissue. Finally, we constructed three candidate EndoMT lineages across multiple subpopulations of ECs and SMCs in human carotid scRNA-seq data (n=46). We examined gene expression over the course of these lineages and identified 73 markers for the presence of EndoMT such as *NRG1* and *DEPP1*.

**Conclusion:** This study reveals the gene expression profile of EndoMT trajectories in human atherosclerotic plaques by combining RNA-seq data from *in vitro* models with single-cell transcriptomic datasets. Our gene expression atlas of EndoMT in atherosclerosis could serve as a reference for future studies, providing novel inroads to study atherosclerotic mechanisms for the development of novel therapies.

## Introduction

Endothelial cells (ECs) form a thin coat on the vasculature’s innermost layer, providing the barrier between blood and vascular tissue. ECs require dynamic and plastic behavior for adequate adaptation upon a large variety of biochemical and hemodynamic stimuli. Proatherogenic stimuli can induce EC plasticity resulting in endothelial-to-mesenchymal transition (EndoMT), where ECs acquire a myofibroblast-like phenotype and lose their EC characteristics^1^. This transition includes a reduction in endothelial marker genes such as VE-cadherin (*CDH5*) and endothelial nitric oxide synthase (*NOS3*) and an increase in fibronectin (*FN1*), alpha smooth muscle actin (αSMA or *ACTA2*) and N-cadherin (*CDH2*)^2^. EndoMT is an important process during arterial and cardiovascular remodeling and can be beneficial or detrimental depending on context^3,4^. Inflammation-induced endothelial dysfunction is a major player in the onset and development of both, atherosclerosis and EndoMT^5,6^.

An important inflammatory driver of EndoMT is the TGFβ superfamily^7–9^, activating a cascade of signaling pathways that transcriptionally reprogram ECs. Upon stimulation of TGFβ, ECs upregulate transcription factor (TF) families e.g., Snail, ZEB and Twist^10^, initiating the molecular mechanism to mesenchymal transition. Cells gain motility by abolishing adhesion junctions between the surrounding cells, excrete matrix metalloproteinases dissolving the extracellular matrix (ECM), and change from an apical-basal polarity towards front-rear polarity, making the cells migratory and invasive^1,11^. Next, upregulation of mesenchymal markers induces a spindle shape morphology, expression of SMC markers, and enhanced ECM production resulting in local tissue fibrosis^2^. There is a large overlap between epithelial to mesenchymal transition (EMT) and EndoMT, and often the terms are used interchangeably. However, they are distinct in their pathology during atherosclerosis^1^.

Based on our current knowledge, EndoMT plays a role in the onset and progression of atherosclerosis. This is mainly based on the identification of EndoMT in mice models^12,13^ and by *in vitro* cell culture experiments^11^. Various studies have identified EndoMT lineages in RNA-seq data from patients, primarily in cancer and metastasis-oriented research fields^14,15^. However, in human atherosclerosis the identification of EndoMT and underlying molecular processes remains largely undetermined as evidence is mainly based on overlapping endothelial and mesenchymal markers in pathological studies^16^. Single-cell transcriptome analysis have been instrumental in improving our understanding of gene regulatory networks involved in atherosclerosis^16^. Using this technology on human plaques, our group recently identified cell transitions as instrumental in atherosclerosis^16–19^. Here, we combined a reference RNA-seq dataset of human coronary artery endothelial cells (HCAEC) undergoing EndoMT under stimulation of TNFα and TFGβ with EndoMT pseudotime lineages created from Cdh5-Cre^ERT2^ Rosa-eYFP apoE^-/-^ lineaged trace mice scRNA-seq data to identify EndoMT. Next, we extrapolated this to EC and SMC scRNA-seq populations of human atherosclerotic plaques, constructing three possible EndoMT lineages and identifying 73 genes including NRG1 and DEPP1, as potential markers for mid-stage EndoMT. Our gene expression atlas could serve as a reference for future studies, providing the unique opportunity to identify EndoMT signatures in human plaques.

## Methods

### Cell stimulation

Human Coronary Endothelial Cells (HCAEC, Promocell, CAT#C-12221) were cultured until 80% confluency was reached. Next, HCAEC were seeded in a concentration of 13,000 cells/well in a 12 wells plate and incubated for 24 hours at 37°C, 5%CO_2_. The following day HCAEC were treated with human TGFβ2 (10ng/ml, Peprotech CAT#100-35-10µg) and/or human TNFα (10ng/ml, Miltenyi Biotec, CAT#130-095-015) in endothelial basal medium MV (EBM-MV, Promocell, CAT#C-22220) supplemented with 0.5% FBS (Corning BV, CAT#2-079-CV) for 0, 2, 6, 12, 24, 48 and 72 hours. At 48 hours EBM-MV was removed from the cells and fresh EBM-MV with the same stimuli was added. At each time point morphology of the cells was checked and cells were resuspended in 100µl TriPure Isolation Reagent (Roche, CAT#11667157001) for further RNA analysis. A detailed overview of the RNA isolation and qPCR analysis is provided in the supplemental methods. Please see the Major Resources Table in the Supplemental Materials.

### RNA-sequencing

RNA library preparation was performed, adapting the CEL-Seq2 protocol for library preparation^20,21^. The initial reverse-transcription reaction primer was designed as follows: an anchored polyT, a unique 6bp barcode, a unique molecular identifier (UMI) of 6bp, the 5’ Illumina adapter and a T7 promoter. Complementary DNA was used for *in vitro* transcription reaction (AM1334; Thermo-Fisher). The resulting amplified RNA (aRNA) was fragmented and cleaned. RNA yield and quality were checked by Bioanalyzer (Agilent).

cDNA library construction was initiated according to the manufacturer’s protocol, with the addition of randomhexRT primer as a random primer. PCR amplification was performed with Phusion High-Fidelity PCR Master Mix with HF buffer (NEB, MA, USA) and a unique indexed RNA PCR primer (Illumina) per reaction. Library cDNA yield was checked by Qubit fluorometric quantification (Thermo-Fisher) and quality by Bioanalyzer (Agilent). Libraries were sequenced on the Illumina Nextseq2000 platform with paired end, 2 × 50 bp (Utrecht Sequencing Facility).

Upon sequencing, fastq files were de-barcoded and split for forward and reverse reads. The reads were demultiplexed and aligned to human cDNA reference (Ensembl v84) using the BWA (0.7.13) by calling ‘bwa aln’ with settings -B 6 -q 0 -n 0.00 -k 2 -l 200 -t 6 for R1 and -B 0 -q 0 -n 0.04 -k 2 -l 200 -t 6 for R2, ‘bwa sampe’ with settings -n 100 -N 100. Multiple reads mapping to the same gene with the same unique molecular identifier (UMI, 6bp long) were counted as a single read.

Downstream analysis of *in vitro* RNA-seq data was performed using DESeq2^22^ (DESeq2 v1.32.0). Differentially expressed genes (DEGs) were determined between control, TNFα, TGFβ and mixed condition. K-means clustering was performed on DEGs identified from all pairwise comparisons between timepoints. Number of K-means clusters was determined by elbow plot (**Supplemental Figure 1D**) and pathway analysis results (enrichR^23^ v3.0).

### scRNA-seq samples

Atherosclerotic lesions from 46 patients (20 female, 26 male) undergoing a carotid endarterectomy procedure were included in the Athero-Express Biobank Study^24^ (AE) at the University Medical Centre Utrecht (UMCU). All patients provided informed consent. Sample processing for the purpose of single-cell RNA sequencing is described elsewhere^16^. Data was processed using Seurat^25^ (v4.0). In short, doublets were omitted by gating between 500 and 10000 unique reads per cell. Data was corrected for sequencing batches. Clustering was based on 30 principal components at resolution 1.25. Sub Clustering of endothelial cells and smooth muscle cells was achieved by clustering based on 15 principal components at resolution 1.2 (n = 672 cells).

scRNA-seq data from micro dissected advanced brachiocephalic artery (BCA) lesions from endothelial lineage-traced (Cdh5-Cre^ERT2^ Rosa-eYFP apoE^-/-^) 18-week Western diet-fed mice, was obtained from Alencar *et al.*^13^. YFP positive cells representing an endothelial background were used for further experiments (n = 434 cells). Data was processed using Seurat^25^ (Seurat v4.0). Clustering was based on 5 principal components at resolution 1.1. Cell population identities were assigned based on marker gene expression and comparisons with (recent) single-cell literature.

### Pseudotime analysis

Pseudotime analysis of scRNA-seq data from human and murine samples was performed using Slingshot^26^ (v2.0), following the Slingshot vignette. Cells were assigned to the lineage if the weight provided by the algorithm was > 0.5. To correlate gene expression with pseudotime, we calculated module scores for gene sets (Seurat) against the cells on the selected lineage and executed Pearson correlation. Pathway analysis was performed using EnrichR (v3.0)^23^. DEG calculations were performed using Seurat. Gene expression analysis along the trajectories was performed using TradeSeq^27^ (v1.6.0). Gene expression patterns were constructed using 1822/1201 DEGs (Adj. p value < 0.01).

### Plaque Bulk RNA-seq analysis

Differential gene expression analysis was performed using DESeq2^22^ (v1.32.0). Using previously published bulk RNA-seq data^17^. Differences between calculated log2FC expression values between gene categories were determined by the Wilcoxon rank-sum test. The effect size was calculated using Cohen’s d.

### GWAS analysis

GWAS summary statistics for coronary artery disease^28^, coronary artery calcification^29^ and carotid intima-media thickness^30^ were accessed via their respective original publication. Downstream processing is described elsewhere^19^.

### Pathways and gene sets

Pathways and gene sets not derived from the *in vitro* experiments were collected from the GSEA (gene set enrichment analysis) database^31,32^ (last accessed December 2021). Only gene sets for homo sapiens were included. In total, 25 gene sets were found using keywords “EndoMT”, “EndMT”, “EMT” and “mesenchymal”. Sets were compared using a hypergeometric test.

### Data

All data was processed in an R (v4) environment. Custom scripts used for this publication are available on github https://github.com/CirculatoryHealth/EndoMT_in_AE.

## Results

### *In vitro* molecular fingerprint of EndoMT

To start defining the molecular signature of EndoMT, we stimulated human coronary endothelial cells (HCAECs) with TNFα and/or TGFβ (10ng/ml) for 0, 2, 6, 12, 24, 48 and 72h (**Figure 1B**). We confirmed the changing morphology using bright-field microscopy and the expression changes of endothelial and mesenchymal marker genes using qPCR (**Supplemental Figure 1B**). After 72h of stimulation, HCAECs had acquired an elongated, spindle-like shape compared to unstimulated HCAECs (representative images **Figure 1C**). The stimulated cells had an upregulated gene expression (2 - 5-fold difference) of mesenchymal markers, such as *VIM*, *FN1* or *TAGLN* and downregulation (0.25 −0.50-fold difference) of EC markers, such as *vWF* and *CD31*, indicative of the mesenchymal transition (**Supplemental Figure 1A**). Although TNFα and TGFβ exposures elicit activation of different pathways within the cells, they are both capable of inducing a mesenchymal phenotype in the HCAEC (**Figure 1D**).

**Figure 1.**
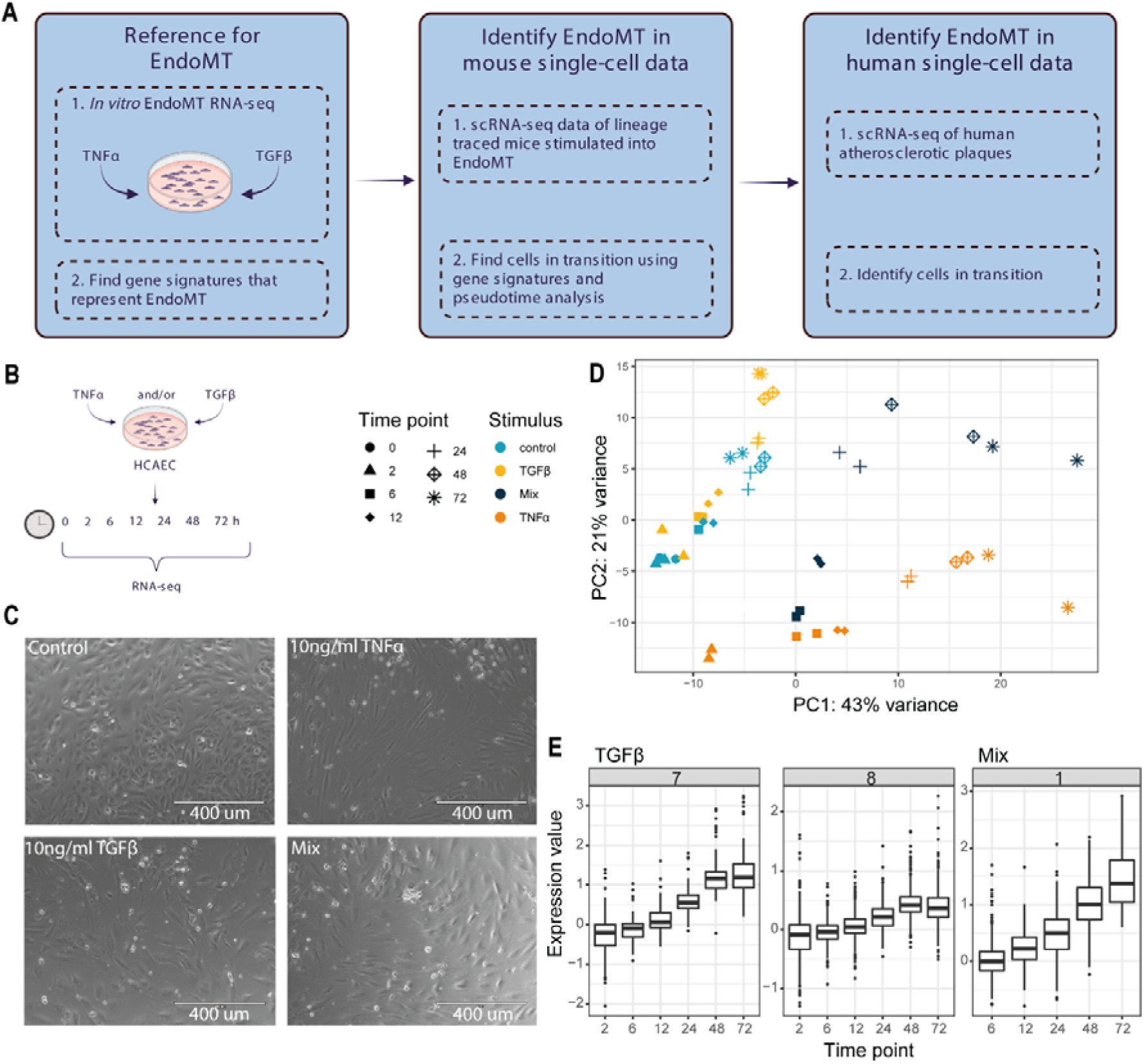
**A**) Visual summary of workflow: stimulated HCAEC identify gene signatures in EndoMT. Next, use Cdh5-Cre^ERT2^ Rosa-eYFP apoE^-/-^ lineage traced mouse scRNA-seq data demonstrate that these signatures can be used to identify EndoMT trajectories in murine plaque tissue. Lastly, similar to the previous, construct candidate EndoMT lineages across multiple subpopulations of ECs and SMCs in human carotid scRNA-seq. **B**) *In vitro* experimental setup: HCAEC are stimulated with 10ng/ml TGFβ2 and/or human TNFα at 7 different time points (0, 2, 6, 12, 24, 48 and 72h) before being processed for RNA-seq **C**) Representative microscopy images of cells unstimulated, stimulated with TGFβ and TNFα. **D**) PCA plot depicting RNA-seq samples from the 4 conditions at different time points. **E**) Expression of K-means genes representative of EndoMT *in vitro* over different time points.

To identify expression patterns that changed over time during EC transition we used RNA-seq data, at time points 0, 2, 6, 12, 24, 48, and 72 h. First, we defined the differentially expressed genes (DEG) between all-time points. With clustering methods (k-means) we were able to identify nine clusters with unique expression patterns per condition over time (TNFα, TGFβ and mix) (**Supplemental Figure 1B**, **Supplemental Figure 2**, **Supplemental Table 1**). To find the gene signatures that represent EndoMT, we selected clusters that showed an increasing gene expression over time and were indicative of EndoMT related processes as determined by pathway analysis (**Supplemental Figure 3**). Based on these clusters in combination with pathway analysis, three unique clusters were selected for further exploration: cluster 7 (n = 69 genes) and 8 (n = 415 genes) from TGFβ stimulated cells, and cluster 1 (n = 141 genes) from cells stimulated with TNFα + TGFβ (“mix”, **Figure 1E, Supplemental Figure 1C**). These three clusters were enriched for GO-terms (top 10) including “extracellular matrix organization”, “endodermal cell differentiation” and “positive regulation of cell migration” (**Supplemental Figure 3**). Cluster 7 included genes such as *SNAI2*, *TPM1* and *TAGLN*. Cluster 8 included *SORT1*, *CNN2* and cluster 1 included *BGN* and *GAS6*.

### Identified gene expression patterns *in vitro* are indicative of EndoMT *in vivo*

To further identify the relevance of our *in vitro* EndoMT signature for transition, we used pseudotime analysis to place individual cells on an inferred time trajectory based on data derived from single-cell -*omics* experiments. We applied this tool to YFP positive cells of scRNA-seq data from plaques derived from Cdh5-Cre^ERT2^ Rosa-eYFP apoE^-/-^ lineage traced mice^13^ (**Figure 2A**). These mice were treated with IL1β to induce EndoMT *in vivo*. The main advantage of using this mouse model is that we were able to trace the cells over time *in vivo* with the certainty of their EC origin. Combining this advantage with the pseudotime inference tool provides a good validation dataset for our in vitro derived clusters.

**Figure 2.**
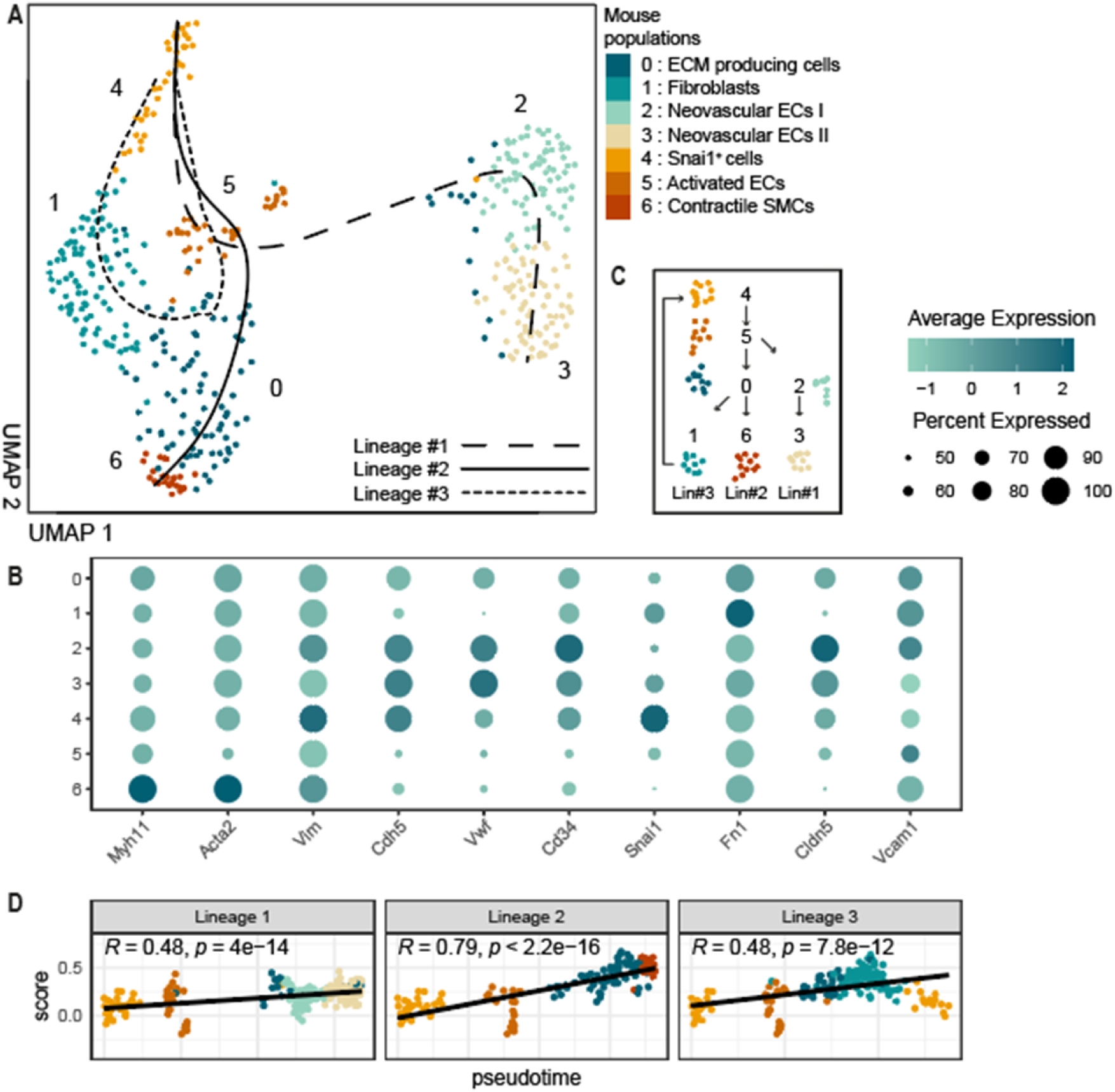
**A**) UMAP visualization of 7 distinct populations derived from Cdh5-Cre^ERT^^2^ Rosa-eYFP apoE^-/-^ lineaged traced mice, including visualization of pseudotime lineages. **B**) Plot showing expression of mesenchymal, endothelial and EndoMT markers across mouse populations 0 to 6 in **A**. **C**) Schematic of 3 putative EndoMT pseudotime lineages shown in **A**. **D**) Depiction of the per cell score calculated from the expression of genes from *in vitro* K-means group TGFβ 7 across pseudotime lineages in **A**.

From the mouse scRNA-seq data, seven sub-populations of ECs and SMCs were identified across 434 cells. Cell populations remained positive for *CDH5*, but during differentiation gained expression of mesenchymal markers such as *ACTA2,* confirming that a transcriptomic phenotypic change was achieved (**Figure 2B**). Cell population 0 was classified as ECM producing cells. Population 1 was positive for fibroblast markers such as *FN1* ^33,34^. Cells from population 2 and 3 show characteristics of neovascular ECs and EC population 4 was *Snai1^+^*. Population 5 showed signs of EC activation and population 6 was positive for contractile SMC markers *MYH11* and *ACTA2*.

Applying the pseudotime analysis algorithm to these populations identified three lineages (**Figure 2A, C**), all originating from *Snai1^+^* cells, moving through cells from the activated EC population and ECM producing cell populations. Lineage #1 tracks along the neovascular ECs and lineage #2 ended in the contractile SMCs. Lineage #3 went through the fibroblasts and circled back to cells from the *Snai1^+^*population.

To validate the three possible EndoMT pseudotime lineages, we calculated a per-cell score for the DEGs present in the *in vitro* EndoMT clusters over time extrapolated the outcome to the 3 lineages. *In vitro* TGFβ cluster 7 was upregulated along the axis of all three lineages (**Figure 2D**). Mix cluster 1 was most upregulated over time in Lineage #2 (**Supplemental Figure 4**). TGFβ cluster 8 was not differentially expressed over time in any of three lineages. Altogether, Lineage #2 is the best matching to the *in vitro* defined EndoMT clusters. From this *in vivo* data, it is suggested that our *in vitro* EndoMT cluster 7 represents a full transition from EC to both a contractile SMC phenotype and an intermediate fibroblast.

### Identifying EndoMT in human atherosclerotic scRNA-seq data

In order to identify genes that associate with EndoMT in human plaques, we proceeded with scRNA-seq data from ECs and SMCs from the Athero-Express biobank^16,24^ (**Figure 3A**, **Supplemental Figure 5A**). 672 cells were divided into eight populations, four with EC identity and four with SMC identity (**Figure 3B**). EC1 is positive for *FN1*, a mesenchymal marker indicative of EndoMT. EC3 showed high expression of *SULF1,* indicating pro-angiogenic endothelial cells^33^. EC2 showed signs of EC activation and EC4 was positive for *ACKR1*, indicating that these cells could be derived from the *vaso vasorum* as neovasculature. SMC1 was identified as myofibroblasts^33^ and SMC2 cells expressed markers indicative of migration. SMC3 was *TBX2* positive^35^ indicating a mesenchymal progenitor cell type and SMC4 showed a contractile SMC phenotype (**Figure 3B, Supplemental Figure 5B**).

**Figure 3.**
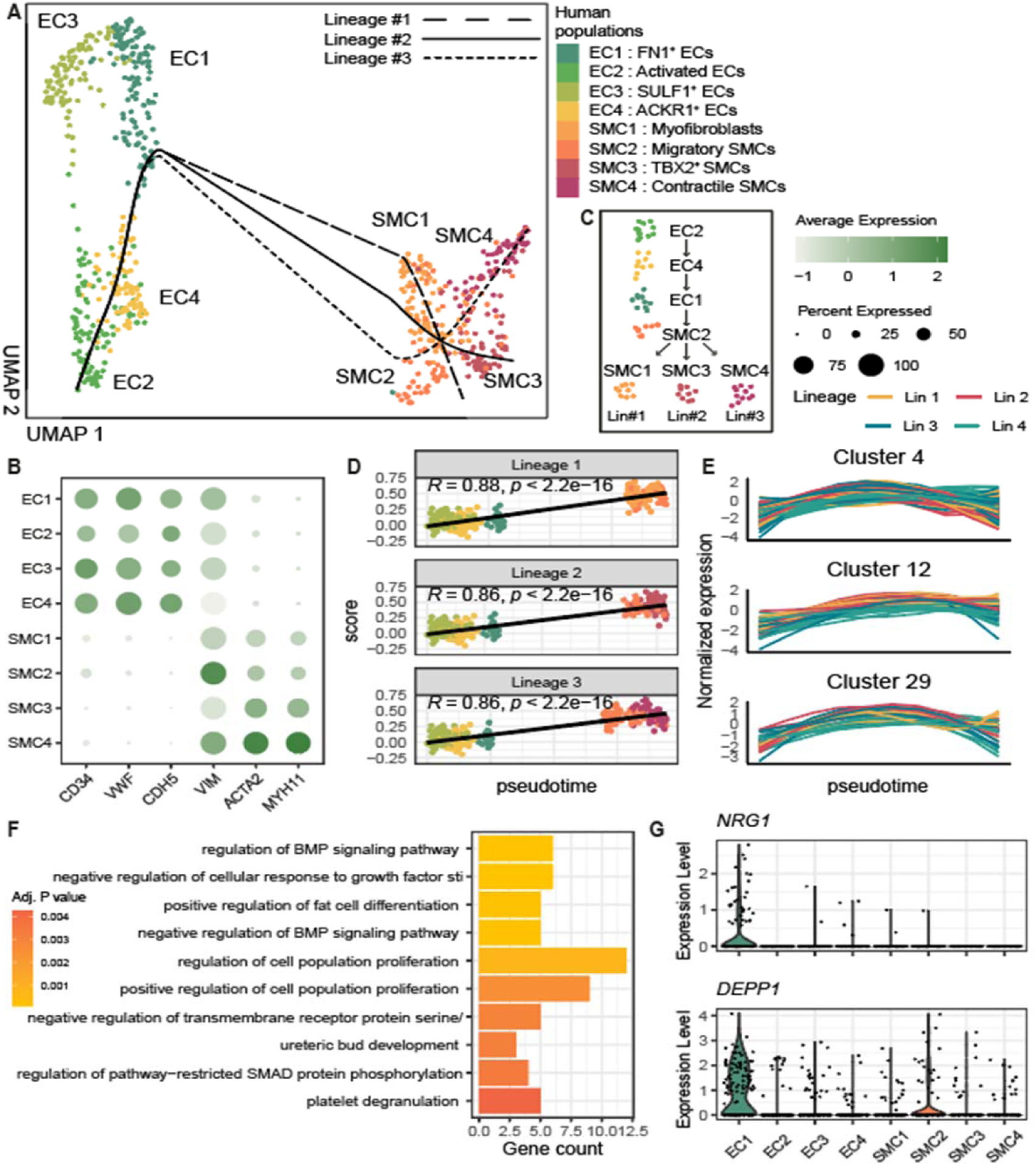
**A**) UMAP visualization of 8 distinct EC and SMC populations derived from scRNA-seq data of human atherosclerotic plaques, including visualization of pseudotime lineages 1, 2 and 3. **B**) Plot showing expression of mesenchymal and endothelial markers across populations from **A**. **C**) Schematic of 3 putative EndoMT pseudotime lineages shown in **A**. **D**) Depiction of the per cell score calculated from the expression of genes from *in vitro* K-means group TGFβ 7 across pseudotime lineages in **A**. **E**) Visualization of the expression DEGs along the lineages. Genes were clustered in 84 clusters based on their changes in expression over pseudotime. The 3 depicted clusters were prioritized based on their involvement in EndoMT (see **F**). Each line represents one gene belonging to the respective lineage. **F**) Pathway analysis results from genes derived from the 3 clusters from **E** (some terms have been truncated). **G**) From the genes depicted in **E**, 2 genes were identified as potential mid-stage markers for EndoMT. Expression levels of potential markers *NRG1* and *DEPP1* in populations of **A**.

To identify possible EndoMT pseudotime lineages we performed pseudotime analysis on this human atherosclerotic data, which resulted in four lineages (**Supplemental Figure 5A**). All lineages originated from EC2, passing via EC4 and EC1 either to SMC2 or EC3. Ultimately three out of four lineages end in one of the remaining SMC populations (lineages #1, #2, and #3) indicative of EndoMT (**Figure 3C**).

We projected the gene expression changes of *in vitro* TGFβ cluster 7 into the three lineages representative of EndoMT (**Figure 3D**, **Supplemental Figure 5C**). Lineages #1, #2 and #3 showed a positive correlation with the *in vitro* EndoMT gene score (R = 0.88, R = 0.86 and R 0.86 resp), indicating that the pseudotime lineages have EndoMT characteristics regardless of the SMC phenotype of the cells at the end of the lineage. The remaining *in vitro* EndMT clusters (Mixed cluster 1 and TGFB cluster 8) also showed a positive correlation with the human pseudotime lineages (Supplemental Figure 5D).

### NRG1 and DEPP1 are associated with mid-stage EndoMT in human atherosclerotic cells

To identify a specific gene expression pattern indicative of active EndoMT in human primary tissue, we examined the gene expression changes of DEGs (adj. p value < 0.01) in the lineages #1, #2 and #3 representing EndoMT characteristics in human atherosclerotic single-cell data. Based on their expression profile across the three lineages, the 1201 DEGs were grouped into 84 clusters (**Supplemental Figure 6**, **Supplemental Table 2**). Within the 84 clusters, various gene expression patterns could be identified over time. For example, clusters 3, 11 (concave expression pattern) and 78 (downwards expression pattern) were enriched for angiogenesis related processes. Cluster 51 showed upwards expression pattern and features genes involved in actin organization.

Next, we examined these 84 clusters to identify possible markers for the active process of EndoMT in human atherosclerosis. Arguably, markers for early and late (end-stage) EndoMT are represented by regular EC and SMC markers. Consequently, we determined for gene expression patterns with upregulation at the mid-stage point across the lineages that are representative for the active ongoing transition. From the 84 clusters, multiple clusters met those criteria showing an upregulation of genes at the mid-stage point across the lineages (**Figure 3E**, **Supplemental Figure 7A**). Clusters 4 (44 genes), 12 (19 genes) and 29 (10 genes) were enriched for GO terms such as “regulation of BMP signaling” (**Figure 3F**, **Supplemental Figure 7A**) and were therefore selected as clusters representing mid-stage EndoMT.

To identify distinct mid-stage EndoMT markers, we narrowed down the 73 mid-stage genes from cluster 4, 12 and 29 **(Supplemental Table 3)** based on their cell type specific gene expression in the scRNA-seq data. *NRG1* and *DEPP1* were identified as mid-stage markers, showing specific expression in SMC2 and/or EC1 cell populations (**Figure 3G**). Additionally, *CTNNAL1* was identified as an early to mid-stage marker, having higher expression in EC4 and EC1 (**Supplemental Figure 7B**).

### Expression of mid-stage EndMT signature genes is affected by plaque morphology and associates with clinical presentation

EndoMT is involved in plaque stability and therefore considered a key contributor to the progression of atherosclerosis^2,12^. To identify associations between the expression of our 73 mid-stage genes representing active EndoMT and plaque composition, we utilized bulk RNA-seq of primary human plaques of 623 patients. The individual EndoMT mid-stage genes were more likely upregulated in plaques with a low macrophage content compared to other genes (Cohen’s d= −0.402, p= 2.8×10^-03^ **Figure 4**). The expression of mid-stage genes was more frequently upregulated in plaques with high collagen (Cohen’s d= 0.781, p= 7.24×10^-8^) and high SMC content (Cohen’s d= 0.823, p=2.21×10^-07^) (**Figure 4**). These results indicate increased activity of EndMT in plaques that contain a high content of SMCs and collagen. Furthermore, two other classical parameters to define plaque composition and vulnerability are calcification (Cohen’s d= 0.266, p= 0.0301) and intra plaque hemorrhage (IPH) (Cohen’s d= −0.278, p= 0.0262). The negative association with the presence of IPH could indicate higher expression of mid-stage genes in a more stable plaque phenotype. Additionally, carotid plaque composition and progression are sex-dependent^36^, we observed a higher expression of mid-stage genes in the females compared to males (Cohen’s d= −0.278, p= 0.06) (**Figure 4**). This could indicate a different contribution of EndoMT to the plaque composition between sexes.

**Figure 4.**
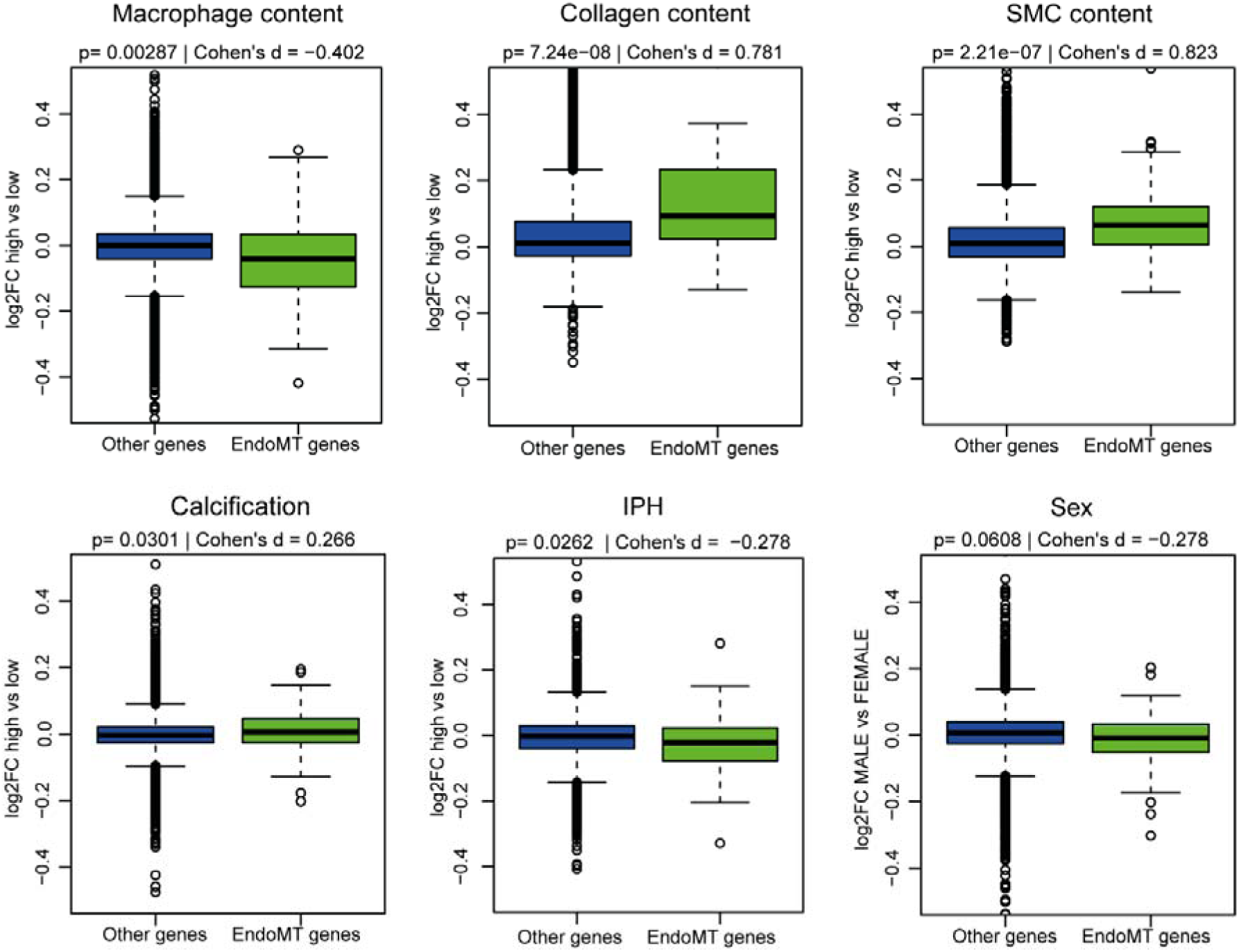
Differential gene expression values among plaques, categorized based on their histological characteristics. Green represent the expression levels of mid-stage EndoMT genes, juxtaposed against the expression of all other genes present in the bulk RNA sequencing data derived from 632 primary carotid plaques. Statistical significance is determined by the Wilcoxon rank-sum test, effect size is determined by Cohen’ s d. IPH; intra plaque hemorrhage, SMC; Smooth muscle cell.

### Markers of mid-stage EndoMT overlap with the GWAS signals for atherosclerotic disease

EndoMT plays a role in the onset and progression of atherosclerosis^1,12^. To study if the identified EndoMT markers overlap with genes within known atherosclerosis-relevant GWAS loci, we took candidate genes for coronary artery disease^28^ (CAD), coronary artery calcification^29^ (CAC) and carotid intima-media thickness^30^ (cIMT) and overlapped this with the 73 mid-stage genes. For CAD three genes were associated with both CAD and EndoMT: *PTGS2*, *FOXC1* and *IGFBP3*. For CAC *CYBRD1* and for cIMT *S100A13*, *MYL12B*, *OAZ1* and *LTBP4* overlapped. These results indicate a possible link with EndoMT and genetic predisposition for atherosclerosis. Enrichment analysis resulted in no significant enrichment for CAD GWAS loci in the EndoMT mid-stage genes.

### Mid-stage EndoMT gene signatures are largely omitted by molecular signatures databases

When identifying EndoMT in scRNA-seq data we often rely on pathway analysis terms that are rooted in cancer research^1^. To determine how representative these gene sets are for human atherosclerosis, we collected 25 pathways and gene sets present in the GSEA database^32^ and compared them to the three mid-stage EndoMT clusters. Sizes of these gene sets ranged from five genes for “Alonso metastasis epithelial to mesenchymal dn” (M3029) and up to 207 genes for “Gotzmann epithelial to mesenchymal transition dn” (M1376) (**Supplemental Figure 7C**).

Overall, there was little overlap between our mid-stage clusters and the 25 pathways and gene sets (**Supplemental Figure 7D**). Similarities between the GO-terms could be explained through overlap of common EC and SMC markers but raises questions on their individual specificity in the context of EndoMT. We noticed overlap between our three clusters and pathways such as “Hallmark epithelial mesenchymal transition” (p value < 0.001, **Supplemental Figure 7D**). Strikingly, Clusters 4 and 29 seem to overlap mostly with genes defined as downregulated, whilst cluster 12 is more associated with sets defined as upregulated genes. Pathway results from cancer research mostly describe end-stage EndoMT, the differences between our three mid-stage clusters and the 25 pathways and gene sets should therefore be carefully considered in the context of atherosclerosis.

## Discussion

We aimed to identify a human EndoMT gene expression signature by combining experimentally induced *in vitro* EndoMT with lineage-traced EndoMT pathways from atherosclerotic mice and extrapolate these results to scRNA-seq data of 46 human coronary plaques. With our *in* vitro model we identified three gene cluster indicative of EndoMT. We used Cdh5-Cre^ERT2^ Rosa-eYFP apoE^-/-^ lineage traced mouse scRNA-seq data to demonstrate that the temporal *in vitro* gene expression changes are reflected in EndoMT trajectories in mice plaque tissue. Finally, pseudotime analysis in subpopulations of ECs and SMCs in human carotid scRNA-seq data (n=46) showed 3 potential EndoMT lineages and 73 possible markers of mid-stage EndoMT. Furthermore, analysis in human carotid bulk RNA-seq data from 641 patients, suggested that higher levels of our 73 mid-stage genes associated with a more fibrous plaque phenotype. Our gene expression atlas of EndoMT could serve as a reference for future studies, providing the unique opportunity to identify EndoMT in human plaques.

To investigate the presence of cells in EndoMT in single-cell data from human carotid artery plaques we used our *in vitro* EndoMT molecular fingerprint. Known gene sets and pathways present in the major pathway analysis tools provided incomplete information on the gene expression over time, especially on early EndoMT. In line, most gene sets contained fibrosis genes, which only represent the end phase of EndoMT. For example, recent attempts at providing an *in vivo* reference EndoMT in human umbilical vein endothelial cells and arterial human pulmonary artery endothelial cells in a hypertension model only provides information at the seven-day time point^37^. Other attempts from the oncology field have shown the importance of temporal data by providing extensive scRNA-seq profiling of EMT in four different cancer cell lines with multiple timepoints between 8 hours and 1 week^38^. From our mid-stage gene clusters, we found that cluster 4 and 29 were more often associated with downregulation and cluster 12 with upregulation pathways or gene sets present in the GSEA database (Supplemental Figure 7D). This indicates that with our approach we were able to detect more subtle mechanisms within the transition process. For our *in vitro* gene set, we used human coronary endothelial cells, relevant to study atherosclerosis. We sequenced the cells at multiple timepoints over the course of 72 hours enabling us to capture early and late gene expression changes over time upon stimulation.

The Cdh5-Cre^ERT2^ Rosa-eYFP apoE^-/-^ lineage traced mouse model allows to study cells that undergo a transcriptomic transition from EC to mesenchymal-like cells with the knowledge that all cells have EC origin. We show the presence of EC derived cells with mesenchymal identity (fibroblasts and contractile SMCs) and intermediate-state cells (*Snai^+^* cells and ECM producing cells) (**Figure 2A, B**) in concordance with previous findings (summarized elsewhere^11^). Even if the fraction of cells actively undergoing transition is small, we confirmed that a phenotypic transition from EC to contractile SMC or fibroblast state is possible in plaque cells over a relatively short time.

From the mouse data, the unidirectionality of the phenotypic switch from EC to mesenchymal is certain, but this does not exclude the possibility of a reversible process under normal physiological conditions. Additionally, from angiogenesis research it is known that partial and reversible EndoMT does exist due to incomplete activation and/or progression of the core EndoMT program^39^. Both a transverse process of EndoMT as well as reversible EndoMT are normal physiological processes initiated by changing conditions in the cellular microenvironment. Similarly, when we consider the transition in mouse lineage #1 (from EC to EC) the structure of the lineage could also be considered as mirrored. In contrast to murine data, we do not have the advantage of knowing the origin of cells in transition in the human data. However, there are large similarities between mice and human cell populations (**Supplemental Figure 8**).

The pseudotime lineage tool uses the populations from scRNA-seq data to construct lineages that follow subtle changes across cells^26^. As a result, cells are assigned to one or multiple lineages that represent different processes. Identifying EndoMT cells solely on marker gene expression is a challenging task since it is a smooth transition across cellular states. Not all known markers are detected in human scRNA-seq (**Figure 2B**, **Supplemental Figure 5B**) and the number of cells actively undergoing transition is low. Additionally, each of these genes has their own unique expression pattern over the dynamic course of EndoMT. Here we have defined three gene clusters spanning 73 genes that represent mid-stage EndoMT genes across human data (**Supplemental Figure 7**). From these 73 genes we identified two markers (*NRG1* and *DEPP1*, **Figure 3G**) that can mark cells actively in transition. These genes have not been implicated in EndoMT previously and could serve useful to identify these mid-stage EndoMT cells in the lesion by pinpointing their location and study the exact cells that are in transition directly.

For this *in vitro* reference set we stimulated endothelial cells in a static condition without taking the blood-vessel structural and cellular components into consideration, which is a limitation of our *in vitro* model. In physiological conditions, EndoMT is influenced by blood flow^40^. Future experiments where the physiological conditions are mimicked would provide more detailed insights in the mechanisms of EndoMT in atherosclerosis. For example, use of 2D and 3D polymeric models to study vascular diseases *in vitro* which incorporate several structural aspects of the vascular wall in non-static conditions^41^, could provide valuable scRNA-seq data. However, it is of high importance that we start with a primary understanding of EndoMT in atherosclerosis.

With our research we identified *NRG1* and *DEPP1* as markers for mid-stage EndoMT. To establish the contribution of these markers to EndoMT in the development of atherosclerosis, further research is needed. We did identify a possible association of higher mid-stage EndMT markers in a stable plaque phenotype. This indicates that the EndoMT markers could be representative of active EndoMT during plaque progression and stability. Nevertheless, the causal contribution of the mid-stage EndoMT genes in EndoMT and plaque composition should be further evaluated. Lastly, we found interesting overlap between the EndoMT markers and the genetic predisposition, however so far it remains unclear how this is involved in EndoMT present in atherosclerosis. With our research we contributed to the identification and primary understanding of the basic characteristics of EndoMT in human atherosclerotic plaque RNA-seq data.

## Supplemental Material

Supplemental Methods

Tables S1–S3

Figure S1-S8

Major Resources Table

---

**Supplemental Figure 1. A)** qPCR relative gene expression scores for *CD31, CDH5, vWF, CDH2, TAGLN, FN1, SNAIL* and *TWIST* after 72h of stimulation with 10ng/ml TGFβ, 10ng/ml TNFα or both (mix) of HCAEC compared to the control condition of 0h. **B**) Elbow plot to determine the number of K-means for K-means analysis. **C**) Graph depicting the number of unique (set size) and overlapping genes (intersection size) per selected K-means group (TGFβ 7, Mix 1, TGFβ 8).

**Supplemental Figure 2.** All K-means gene clusters expression levels, presented per stimulus condition (control, TGFβ, TNFα or mix), per time point (0-72h).

**Supplemental Figure 3**.Pathway analysis results from all genes derived from the K-means clusters from **Supplemental Figure 2**, presented per stimulus condition (control, TGFβ, TNFα or mix).

**Supplemental Figure 4.** Depiction of the per cell score calculated from the expression of genes from *in vitro* K-means group TGFβ 8 (top) and mix 1 (bottom) across pseudotime lineages in mice.

**Supplemental Figure 5. A)** UMAP visualization of 8 distinct populations derived from human atherosclerotic plaques, including visualization of pseudotime lineages 1, 2, 3 and 4. **B**) Plot showing expression of cell population markers and EndoMT markers across populations in A. **C**) Depiction of the per cell score calculated from the expression of genes from *in vitro* K-means group TGFβ 7 across pseudotime lineage 4. **D**) Depiction of the per cell score calculated from the expression of genes from *in vitro* K-means group TGFβ 8 (top) and mix 1 (bottom) across pseudotime lineages in A.

**Supplemental Figure 6.** Visualization of the expression of 84 clusters of DEGs along the 4 human lineages (see **Supplemental Figure 5A**). Genes were clustered based on their changes in expression over pseudotime. Each line represents one gene belonging to the respective lineage.

**Supplemental Figure 7. A)** Pathway analysis results from 3/84 gene clusters derived from human pseudotime (see **Figure 3E**). **B**) Expression levels of *CTNNAL1* in human scRNA-seq populations. **C**) Number of genes per geneset from **A** and pathways or genesets selected from the GSEA database. **D**) Heatmap showing overlap between pathways or genesets (hypergeometric test).

**Supplemental Figure 8.** The expression of human population specific DEGs across the mouse populations represented by a per cell score. Numbers indicate the amount of traceable human DEGs in mouse data.

